# Comparative analysis of microbial diversity across temperature gradients in hot springs from Yellowstone and Iceland

**DOI:** 10.1101/841700

**Authors:** Peter T. Podar, Zamin Yang, Snædís H. Björnsdóttir, Mircea Podar

## Abstract

Geothermal hot springs are a natural setting to study microbial adaptation to a wide range of temperatures reaching up to boiling. Temperature gradients lead to distinct microbial communities that inhabit their optimum niches. We sampled three distant but chemically similar hot springs in Yellowstone and Iceland that had outflows and presented a wide range of temperatures. The microbial composition at different niches was determined by deep DNA sequencing of rRNA gene amplicons. Over three dozen phyla of Archaea and Bacteria were identified, representing over 1700 unique organisms. We observed a significant reduction in the number of microbial taxa as the temperature increased from warm (38°C) to boiling. The community structure was primarily driven by temperature and, at high taxonomic levels, was similar between Yellowstone and Iceland, suggesting that the environment rather than biogeographic location play the largest role in the microbial ecology of hot springs. We observed significant endemism at genus level, especially in thermophilic phototrophs, potentially driven by distinct environmental conditions and dispersal limitations.

## Introduction

Archaea and Bacteria inhabit every environment on Earth, including many that are inhospitable to multicellular life, such as hot springs. As geothermal water cools, outflowing from the source, temperature, chemical and redox gradients form. Distinct microbial communities occupy the various niches of such gradients, based on individual species adaptation to different temperature and chemical optima [1,2]. Hot springs around the world have been used as natural laboratories to study the effect of environmental parameters on microbial evolution, diversity and physiology [3–7]. Extreme temperature and pH values have been shown to have the largest contribution in restricting microbial diversity, although the effects were not linear and were dependent on the system and impacted by other environmental parameters [8–10].

Hot springs have also been used to test hypotheses on factors and mechanisms that lead to microbial diversification and biodiversity patterns [11]. The hypothesis that microbes in general have a high dispersal rate, that would homogenize genetic variations that may arise as result of local ecological and evolutionary events, has been challenged by studies of microbes in geothermal hot springs. Because thermophilic microbes are unlikely to survive too long outside of their habitat, they have limited dispersal ability over large geographical distances (thousands of kilometers). Therefore, while microbial communities that inhabit geochemically similar hot springs on different continents are expected to be physiologically and taxonomically similar, some of the individual species may evolve as endemic populations, similar to plants and animals on distant islands. This has been demonstrated for thermophilic *Synechococcus* (Bacteria) and *Sulfolobus* (Archaea) [12,13].

Here we studied the microbial diversity across a temperature gradient in three alkaline hot springs from Yellowstone National Park (YNP) and Iceland that have not been previously characterized microbiologically. We hypothesized that even though the individual hot springs are geographically isolated, they would share the same general microbial community composition (and potentially physiological activities) at similar temperatures along the gradient. Also, because the graphical distance separating the hot springs in North America from those in Iceland, we also aimed to detect genetic variation between the shared taxa at matched temperatures.

## Materials and Methods

### Icelandic hot spring samples

Microbial mats, sediments, and water samples were collected on June 9, 2016 at a hot springs field in the village of Flúðir, Iceland (GPS coordinates 64º08’13” N 20º18’34” W). The main hot spring, Vaðmálahver, is alkaline (pH ~8.5) and the source water is 98°C (boiling). The outflow of Vaðmálahver gradually cools and the water discharges in a nearby river. Several hot spring sources from the same site discharge in Hverahólmi, the oldest public swimming lagoon in Iceland (1891). The temperature in the main source as well as in the outflow, sediments and in the microbial mats was measured using a Fisher stainless steel temperature probe at the end of a long (10 ft) wire cable positioned either manually or with a telescopic pole. Water and sediment gravel from the main source were collected using a stainless-steel cup at the end of a telescopic pole and immediately poured into sterile 100mL Pyrex glass bottles, capped with no air headspace and secured using butyl rubber stoppers and aluminum crimps. Microbial mats and sediments (~1-2 grams) were collected using sterile syringes and stainless-steel spatulas and placed into plastic tubes containing ceramic beads and 750µl Xpedition™ Lysis/Stabilization Solution (Zymo Research, Irvine, CA) and lysed by bead-beating for one minute with a battery-operated tube shaker. That ensured cellular lysis, inactivation of degradative enzymes and stabilization of the DNA until further processing. A total of seven different spots were sampled from and around the Vaðmálahver spring, ranging from 98 to 47°C. (**Figure 1 and Table 1**). The main source of an adjacent spring (temperature of 92°C) that flows into Hveraholmi was also collected, as well as mat and water samples from the swimming pool (38°C). The microbial community from the swimming lagoon (250 ml water sample) was collected on a Sterivex 0.2 mm syringe filter and preserved by adding Xpedition solution into the filter cartridge and then capping. A microbial mat sample was collected from a rock in the pool and processed as were the other mat samples. After reaching the laboratory the lysed samples were stored at −20°C until DNA extraction or at 4°C (samples for geochemical analyses).

**Figure 1.**
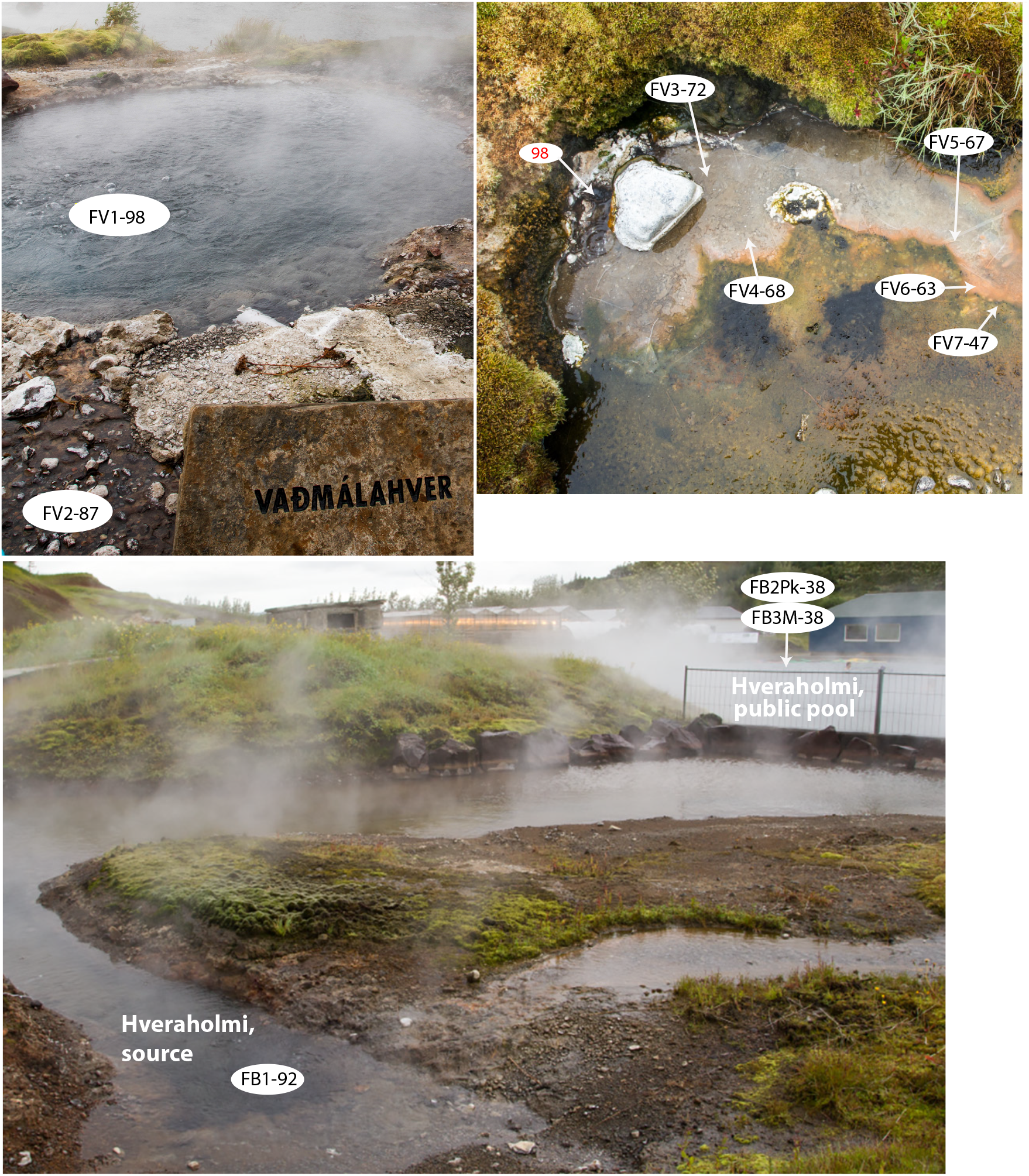
Sampling sites of the hot spring features at Flúðir, Iceland.

**Table 1.** Environmental samples used in the study.

| SampleID | Location | Thermal Feature | Temperature, °C | Replicate |
| --- | --- | --- | --- | --- |
| MP1A | YNP | MirrorPool | 52 | 1 |
| MP1B | YNP | MirrorPool | 52 | 2 |
| MP1C | YNP | MirrorPool | 52 | 3 |
| MP2A | YNP | MirrorPool | 58.6 | 1 |
| MP2B | YNP | MirrorPool | 58.6 | 2 |
| MP2C | YNP | MirrorPool | 58.6 | 3 |
| MP3A | YNP | MirrorPool | 65.5 | 1 |
| MP3B | YNP | MirrorPool | 65.5 | 2 |
| MP3C | YNP | MirrorPool | 65.5 | 3 |
| MP4A | YNP | MirrorPool | 69 | 1 |
| MP4B | YNP | MirrorPool | 69 | 2 |
| MP4C | YNP | MirrorPool | 69 | 3 |
| MP5A | YNP | MirrorPool | 72.5 | 1 |
| MP5B | YNP | MirrorPool | 72.5 | 2 |
| MP5C | YNP | MirrorPool | 72.5 | 3 |
| MP6A | YNP | MirrorPool | 78 | 1 |
| MP6B | YNP | MirrorPool | 78 | 2 |
| MP6C | YNP | MirrorPool | 78 | 3 |
| MP6D | YNP | MirrorPool | 83 | 1 |
| FV2 | Iceland | Vaðmálahver | 87 | 1 |
| FB1 | Iceland | Hverahólmi sr. | 92 | 1 |
| FB2.Pk | Iceland | Hverahólmi | 38 | 1 |
| FV3.Mat | Iceland | Hverahólmi | 38 | 1 |
| FV1 | Iceland | Vaðmálahver | 98 | 1 |
| FV3 | Iceland | Vaðmálahver | 72 | 1 |
| FV4 | Iceland | Vaðmálahver | 68 | 1 |
| FV5 | Iceland | Vaðmálahver | 67 | 1 |
| FV6 | Iceland | Vaðmálahver | 63 | 1 |
| FV7 | Iceland | Vaðmálahver | 47 | 1 |
| HD1 | Iceland | Hurðarbak | 100 | 1 |
| HD2 | Iceland | Hurðarbak | 81 | 1 |
| HD3 | Iceland | Hurðarbak | 67 | 1 |
| HD4 | Iceland | Hurðarbak | 61.4 | 1 |
| HD5 | Iceland | Hurðarbak | 45.5 | 1 |

The second location in Iceland was also an alkaline spring (pH 8.0) at Hurðarbak, in the Borgarfjörður region [GPS coordinates 64º41’18” N, 21º24’10” W] approximately 80 km NW of Flúðir. Samples were collected on June 16, 2016. Five sampling spots were selected, with temperatures ranging from 99-102°C (the source spring) to 46°C in the outflowing channel (**Figure 2** and **Table 1**). Sample collection and processing were performed as described above.

**Figure 2.**
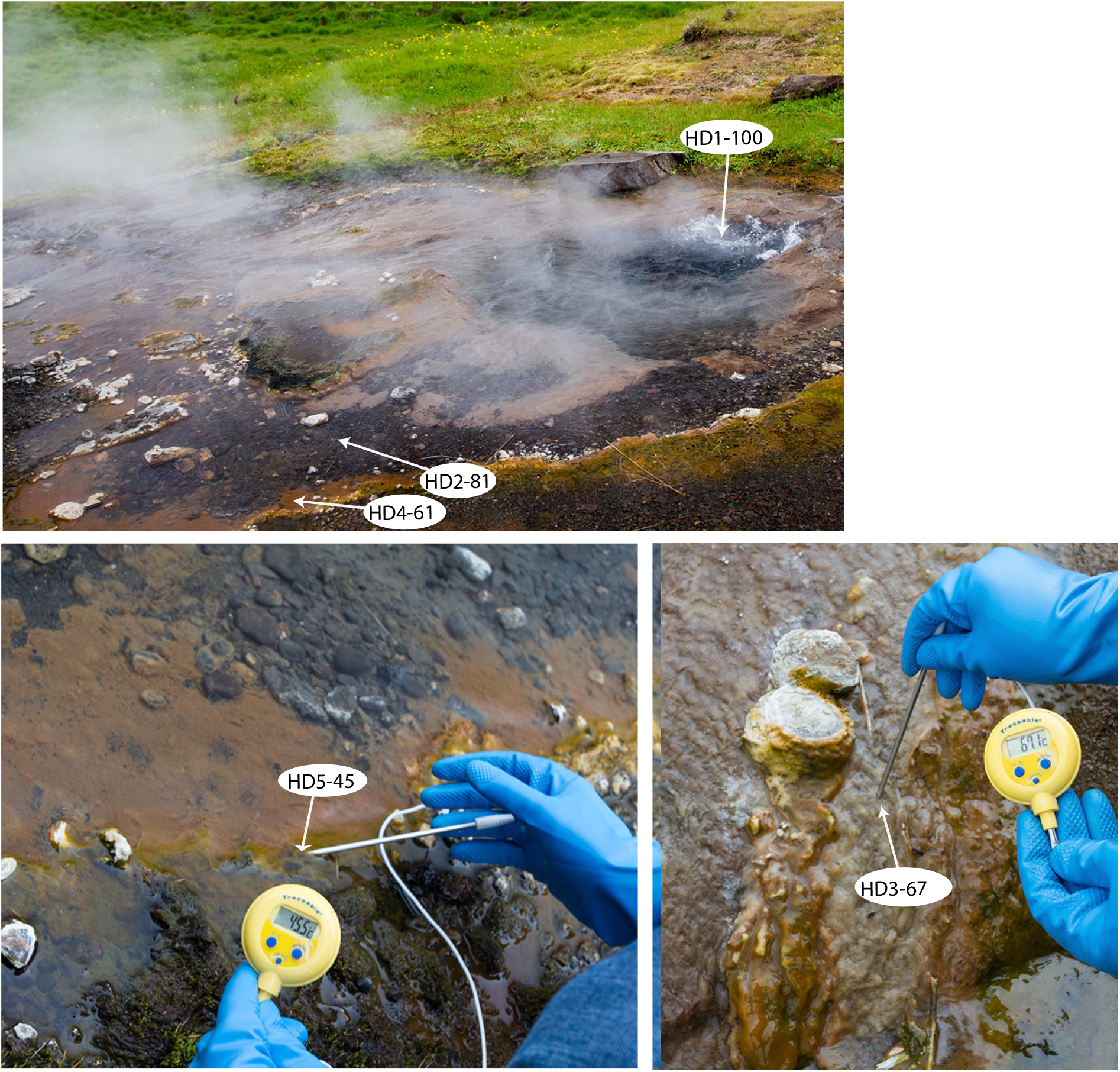
Sampling sites from the hot spring at Hurðarbak, Iceland.

### Yellowstone National Park hot spring samples

Microbial mats, sediment, and water samples were collected on December 31, 2016 at Mirror Pool, an alkaline (pH 8.0) thermal feature from the Upper Geyser Basin [GPS coordinates 44º28’59” N, 110º51’01” W]. Nine sites were sampled along the main pool and gradient outflow of the spring, where temperatures ranged between 83°C-52°C (**Figure 3** and **Table 1**). Because the outflow of the pool was larger than for the hot springs in Iceland it was feasible to identify spots with the same temperature. Therefore, three adjacent replicate samples separated by less than 10 centimeters were collected for each temperature value, to determine the degree of diversity fluctuation across small scales. The samples were collected and processed as described above.

**Figure 3.**
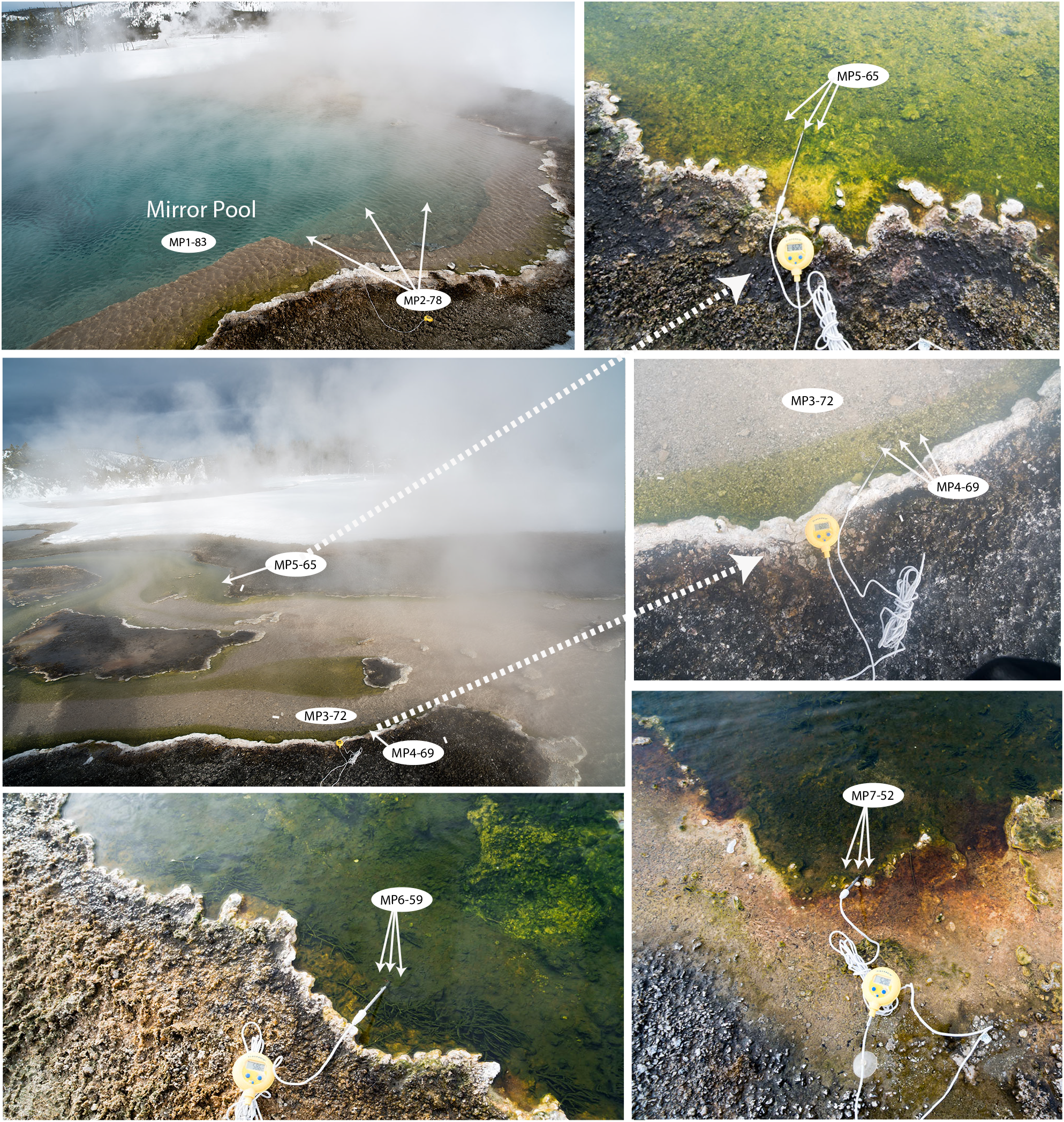
Sampling sites of Mirror Pool, Yellowstone National Park.

### Water chemistry analysis

The chemical composition of the hot spring water samples was performed at the University of Tennessee Knoxville Water Quality Core Facility. Carbon and nitrogen content were determined by thermal combustion and infrared detection with a Shimadzu carbon/nitrogen analyzer. The concentration of metals was measured by inductively coupled argon plasma (ICP) optical emission spectrometry using a Thermo-Scientific iCAP 7400 ICP spectrometer. Ions were measured by ion chromatography with a Thermo-Scientific®/Dionex ICS-2100 (anions) and ICS-1100 (cations), with background suppression for low detection limits.

### DNA extraction

Total genomic DNA from environmental samples was isolated using the ZR Soil Microbe DNA Kit (Zymo Research) following the manufacturer’s protocol. To isolate DNA from the high temperature sediment samples collected in the larger volume bottles, 25 mL of those samples subjected to centrifugation (12,000 × *g* for 20 minutes), the water carefully decanted, and the pellet suspended and lysed using the Zymo Lysis/Stabilization Solution following by processing as above. The concentration of DNA was determined using a Qubit dsDNA HS assay kit and fluorometer (Thermo Fisher Scientific).

### Microbial SSU rRNA gene amplicon sequencing

The V4 hypervariable region of the small subunit ribosomal RNA gene (SSU rRNA) was amplified using universal bacterial/archaeal 515F and 806R primers fused to Illumina sequencing adapters, following the procedure developed by Lundberg et al [14]. To increase the coverage of archaeal groups not recognized effectively by the standard 515F-806R primers (5’ GTGCCAGCMGCCGCGGTAA and 5’ GGACTACHVGGGTWTCTAA, respectively), we supplemented the reaction with modified versions that included 20% 515FCren (5’ GTGKCAGCMGCCGCGGTAA, for Crenarchaeota), 5% 515FNano (5’ GTGGCAGYCGCCRCGGKAA, for Nanoarchaeota) and 5% 805RNano (5’ GGAMTACHGGGGTCTCTAAT, for Nanoarchaeota), similar to what was described in [15]. 12-nucleotide barcode sequences were incorporated into the second stage amplification reaction to enable sample multiplexing. The final amplicons were pooled, purified using Agencourt Ampure XP bead and quantified using Qubit. A diluted purified pooled amplicon sample (9 pM), containing 20% phiX DNA was denatured and sequenced (2 × 250 nt) on an Illumina MiSeq instrument (Illumina Inc, San Diego, CA) using a v2 500 cycle kit, according to manufacturer’s protocol.

### Amplicon sequence analyses

The amplicon primer regions were trimmed from the raw fastq sequence files using cutadapt [16]. The sequence reads were then de-multiplexed based on barcode sequences using the QIIME [17] python script split_libraries_fastq.py followed by splitting by individual samples using split_sequence_file_on_sample_ids.py. Demultiplexed fastq paired reads were imported into QIIME2 v2019.7 [18] on a desktop computer. The reads were paired with vsearch [19], quality filtered and denoised using Deblur [20]. Resulting amplicon sequence variants (ASV) were aligned and used to generate a phylogenetic tree using the align-to-tree-mafft-fasttree pipeline from the q2-phylogeny plugin. To calculate alpha-diversity metrics (observed OTUs, Pielou’s eveness, Shannon’s index and Faith’s Phylogenetic Diversity [21]), beta diversity metrics (weighted UniFrac [22] and Bray-Curtis dissimilarity), and input the resulting matrices into principle coordinate analyses (PCoA) and visualization plots, we used the q2-diversity workflow, with rarefaction to 5000 sequences per sample (based on plateauing of the observed diversity and retaining of all samples). A general temperature classification of samples was generated by assigning each sample to groups separated by 5 °C (from 40 °C to 100 °C). Environmental parameters that could impact alpha diversity were tested using Sperman correlation and analysis of variance (ANOVA), using q2 diversity alpha-correlation and q2 longitudinal [23]. To test for factors that contribute to microbial diversity differences between the samples (actual temperature, general temperature, location, hot spring) we used multi-way permutational multivariate analysis of variance (PERMANOVA) (q2 diversity adonis tests), comparing the variance explained by the various parameters singly or in combinations. Pairwise tests within metadata categories were performed by one-way PERMANOVA using the q2 diversity beta-group-significance.

To assign taxonomy to ASVs we used the q2-feature-classifier [24] (classify-sklearn) against the Silva-132-99 SSU rRNA database. The scripts and parameters are provided as a supplementary file. The raw fastq files were submitted to the NCBI SRA.

### Phylogenetic analyses

Phylogenetic trees to compare selected Yellowstone and Iceland ASVs with related organisms from GenBank were generated using PhyML in the software package Geneious (www.geneious.com). Blastn search algorithm was used to identify relatives of the Yellowstone and Iceland bacteria and archaea in public sequence databases.

## Results and Discussion

### Geochemical comparisons of the three hot springs

The three hot springs were selected because of their high temperature (80-100°C at the source), similarities in pH and the presence of discharge channels with gradual cooling that harbor distinct microbial mats (**Figures 1-3**). Mirror Pool is a large (~15×20 meters) non-erupting, deep, light blue color pool, in the Upper Geyser Basin thermal region (Cascade Group complex) of Yellowstone National Park. Abundant silica deposits are present both in the pool and on its edges and the outflow channel. The temperature and pH we recorded are very similar to those reported in the YNP Research Coordination Network database (http://rcn.montana.edu/Features/Detail.aspx?id=8999), 76-80°C and pH 8, measured in 1999. Similar to other alkaline-siliceous chloride-type springs in that thermal region [25], Mirror Pool has relatively high levels of chloride, sodium, silica and arsenic but is low on sulfur or sulfate, calcium and magnesium **(Table 2)**. Its low flow rate and close proximity to the forest line are probably linked to the relatively high dissolved organic carbon (~100mg/l). Unlike Mirror Pool, the two sampled hot springs in Iceland were much smaller (<2 m in diameter) and actively discharging, likely explaining their lower dissolved organic carbon content (10-20 mg/l). The overall mineral content of both springs was also lower than that of Mirror Pool, although they had higher levels of sulfate, calcium and iron, as previously reported for Vaðmálahver [26,27].

**Table 2.** Chemical composition of hot spring water samples. All values in mg/l. TC, total carbon; TIC, total inorganic carbon; TOC, total organic carbon. Total nitrogen, nitrite, Mn, Zn, Cd, Co, Ba, Se, Tl, Ag: not detected (<0.01 mg/l).

|  | Mirror Pool | Vaðmálahver | Hurðarbak |
| --- | --- | --- | --- |
| TC | 101.6 | 24.39 | 11.94 |
| TIC | 4.84 | 3.01 | 2.24 |
| TOC | 96.79 | 21.38 | 9.71 |
| Cl | 271.79 | 24.89 | 32.01 |
| NO3 | 0.19 | 0.06 | 0.07 |
| SO4 | 13.78 | 54.97 | 57.13 |
| HPO4 | 0.1 | 0.13 | 0.15 |
| F | 19.34 | 1.14 | 1.84 |
| Na | 400.09 | 78.23 | 69.87 |
| K | 10.3 | 0.73 | 0.83 |
| Mg | 0.02 | 0.11 | 0.08 |
| Ca | 0.75 | 5.89 | 4.26 |
| Al | 0.33 | 0.05 | 0.05 |
| Cu | 0 | 0 | 0.01 |
| Fe | 0.04 | 0.15 | 0.17 |
| Si | 119.55 | 31.38 | 20.23 |
| S | 4.69 | 25.03 | 23.01 |
| Ni | 0.03 | 0.01 | 0 |
| Pb | 0.18 | 0.04 | 0.03 |
| Cr | 0.01 | 0.01 | 0.01 |
| Be | 0.01 | 0 | 0 |
| As | 1.32 | 0.01 | 0.02 |
| Sb | 0.03 | 0 | 0 |
| Y | 0.01 | 0.01 | 0.01 |

### Temperature influences the microbial alpha diversity

The combined sequencing of the SSU rRNA amplicons from all samples resulted in over 5.7 million sequences. After de-multiplexing, quality-based filtering, denoising, chimera and singleton removal, the number of sequences for individual samples ranged from 5,315 to 332,920, with an average of ~115,000 sequences per sample and a total of 1729 amplicon sequence variants (ASVs)(unique taxa). Because the type of clustering algorithm and selection of similarity level impacts the number of traditional operational taxonomic units (OTUs), we only used ASVs for calculation of diversity indices.

For studying the microbial diversity within each sample (alpha diversity) we used both direct counts of the number of ASVs as well as metrics that take into account the evenness of species richness (Pielou’s evenness), abundance and distribution of the taxa (Shannon’s index) or the phylogenetic diversity (Faith’s diversity). In Mirror Pool, where we were able to take spatially separated samples at the same temperature, alpha diversity was highly similar between replicates (±0.2-12% standard deviation). Temperature had the largest impact and was inversely correlated with the number of detected taxa (ASVs) and phylogenetic diversity (Spearman p=0.0063 and p=0.0000, respectively) (**Figure 4 and 5**), but did not significantly correlate with evenness or abundance and distribution. The decline in number and diversity of microbial taxa with temperature does not appear to be linear and is steeper in the ranges corresponding to the transition between mesophily and thermophily (35-45 °C) and between thermophily and hyperthermophily (>80 °C). Such non-linear relationships have been previously reported for hot spring communities in Canada, New Zeeland, US (Nevada) and Thailand where wide temperature ranges were present within individual thermal systems [28,10,8]. ANOVA tests of potential multiple effects on the alpha diversity confirmed that, while temperature had the largest influence (P-Value from F-Ratio=3.4e-05), the individual thermal feature was significant too (P-Value from F-Ratio=3.6e-03, passing pairwise T-tests with BH-FDR). When only strict thermophilic temperature values were considered (50-80 °C, the range sampled in Mirror Pool), the effect on alpha diversity was absent, which may explain reports of no temperature effect on species richness [29]. Also, when analyzing the temperature distribution of non-phylogenetic alpha diversity indices (Pielou’s, Shannon’s), we observed that samples spanning the 67-80 °C were sharply elevated relative to what appears to be linear distribution across the other two temperature ranges (**Figure 4**). Because in the 67-80 °C range there are major shifts in the microbial communities, with the dominant photosynthetic groups (Cyanobacteria, Chloroflexi) being replaced by various extreme thermophilic taxa (Thermi, Aquificae, Crenarchaeota), we calculated alpha diversity after removal of ASVs assigned to the photosynthetic bacteria. The split of the non-phylogenetic indices across temperature ranges was reduced, with essentially no effect on temperature correlation for raw ASVs and Faith’s phylogenetic diversity (**Figure 4**). These results highlight the importance of sampling multiple, narrow temperature ranges in hot spring environmental sampling and the choice of diversity indices in studying such extreme environments.

**Figure 4.**
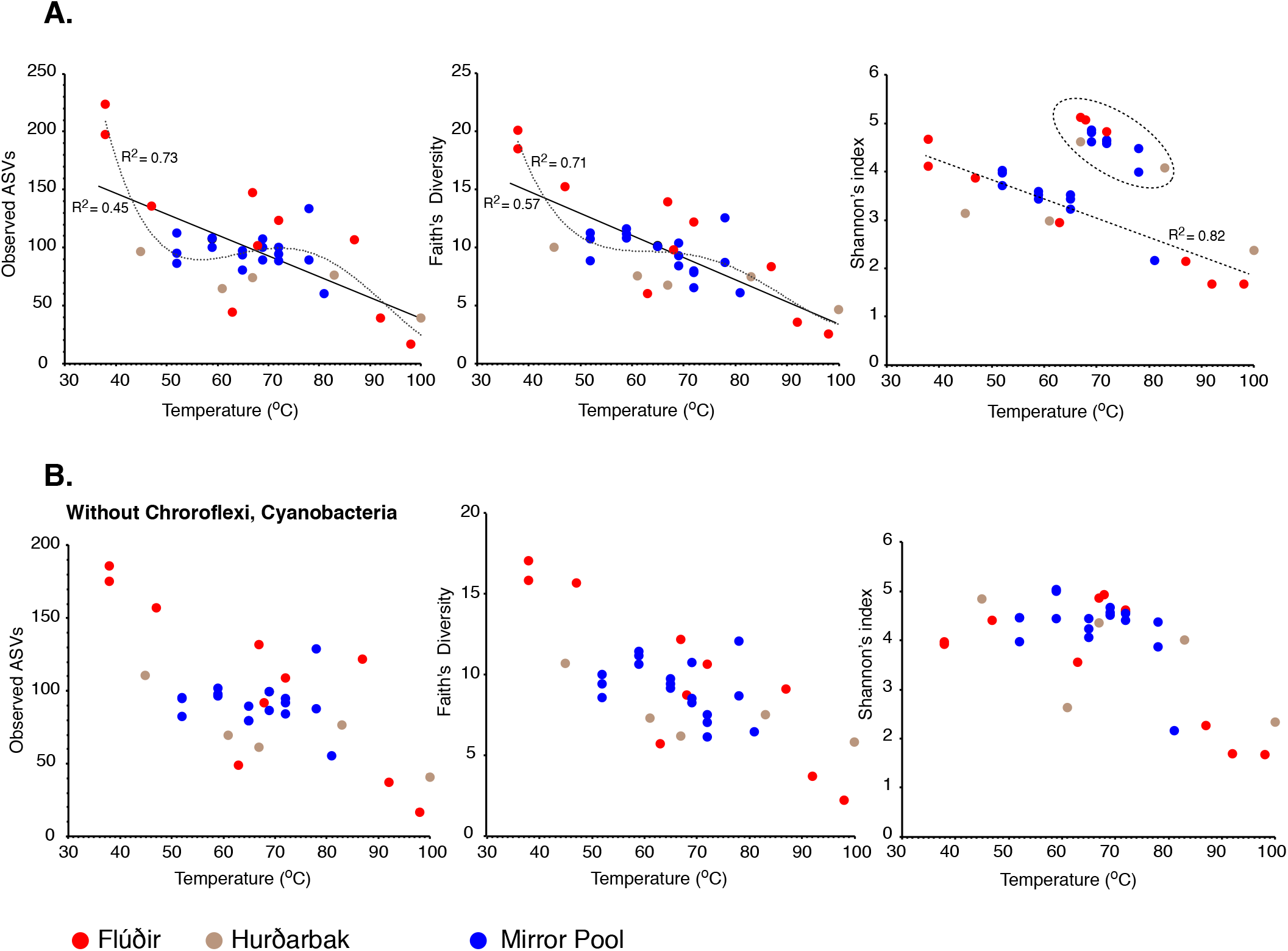
Microbial alpha diversity versus temperature. **A.** Scatterplots of various diversity indeces versus temperature. Linear or polynomical regression and goodness of fit are shown. For the Shannon’s index, the effect of excluding the upper cluster values (circled) on linear regression fit is shown. **B.** Scatterplots of diversity indeces versus temperature after excluding the Cyanobacteria and Chloroflexi sequences.

**Figure 5.**
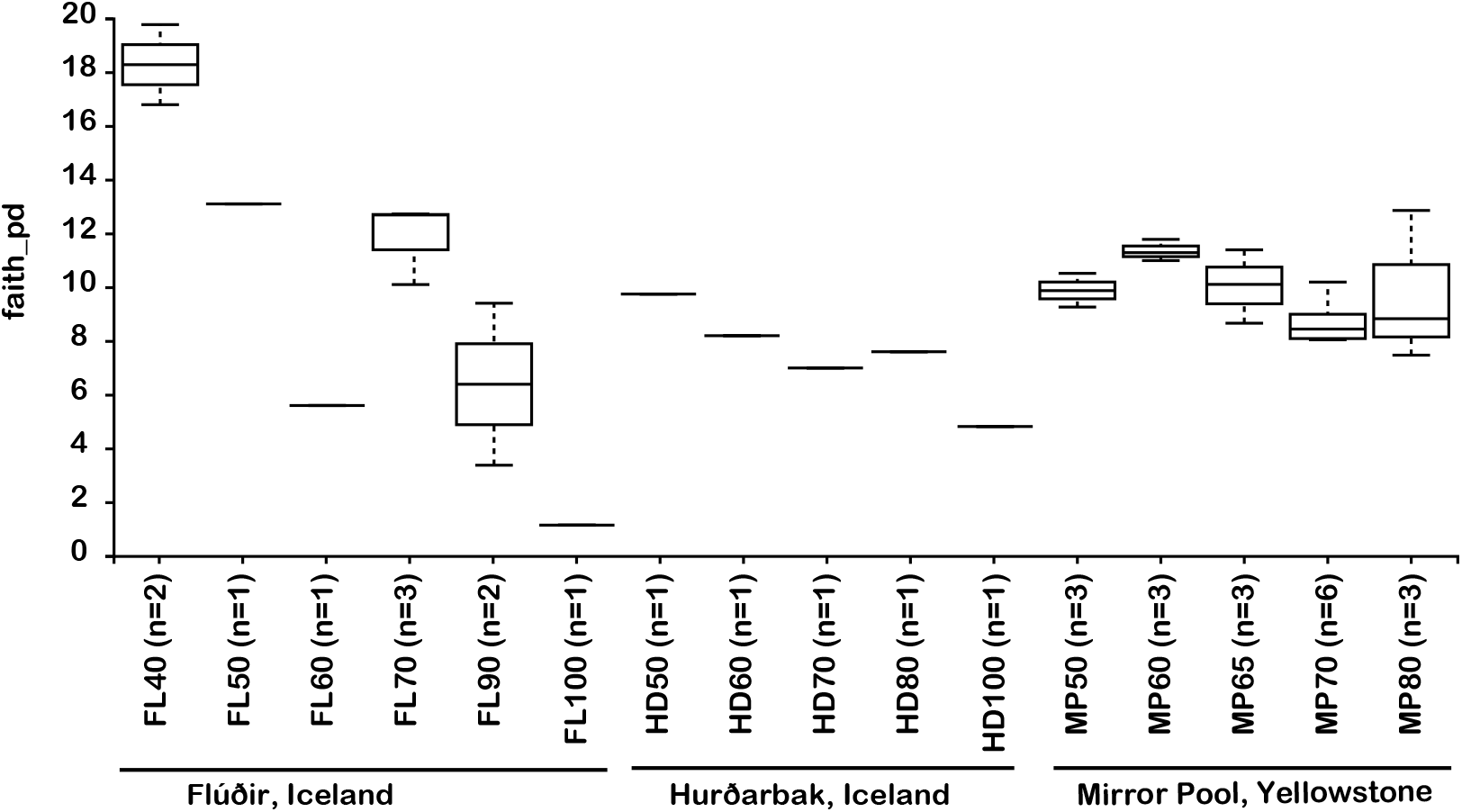
Alpha diversity correlation to temperature. Faith’s phylogenetic distance was tested relative to temperature groups for the three hot spring locations (Kruskal-Wallis test p=0.03).

### Temperature and biogeography effects on beta diversity

We next inspected the structure of the communities across temperatures and hot springs using projections of similarity matrices in principal coordinates space. The weighted unifrac matrix explained most of the variation in the three main coordinates, ~77%. General temperature was the most important factor (PERMANOVA pseudoF=10.4, p=0.001, ANOSIM R=0.77, p=0.001), which is also evident on the based on the emperor PCoA plot (**Figure 6 and 7**), where samples are distributed primarily by temperature rather than location or thermal feature. Both location (Iceland or YNP) and individual thermal features had smaller but statistically significant contributions, probably reflecting both specific taxa and temperatures. A test of the combined effect of temperature and location to beta diversity revealed over 80% of the variation explained by those factors (ADONIS R^2^=0.877, p=0.001). To our knowledge this is the first time that the structure of microbial communities across temperature gradients from several geographically isolated locations were found to be primarily linked to specific temperatures rather than biogeography and geochemistry.

**Figure 6.**
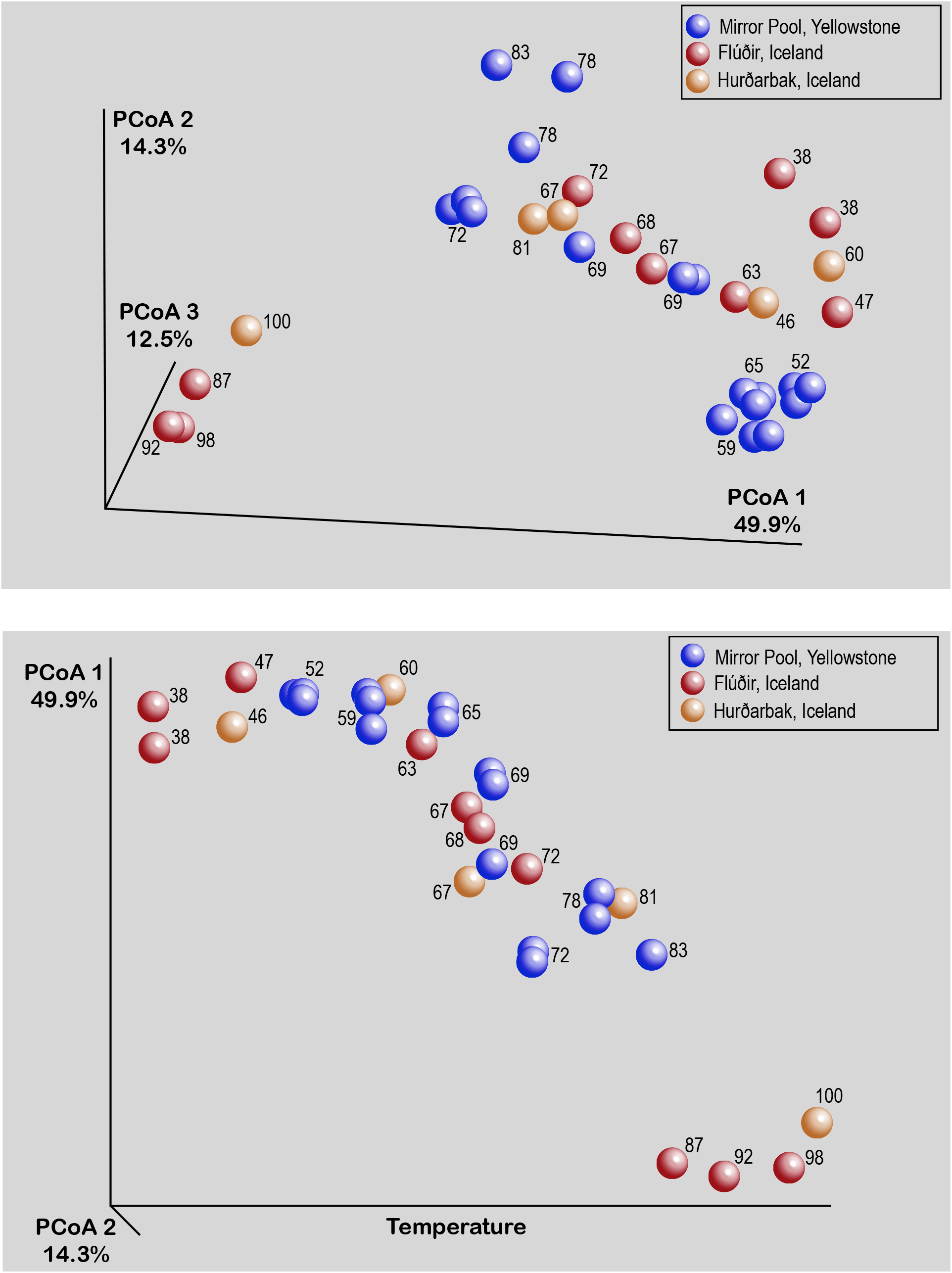
Representation of hot springs microbial beta diversity through EMPeror plots of the principal coordinates analysis output for weighten unifrac distances.

**Figure 7.**
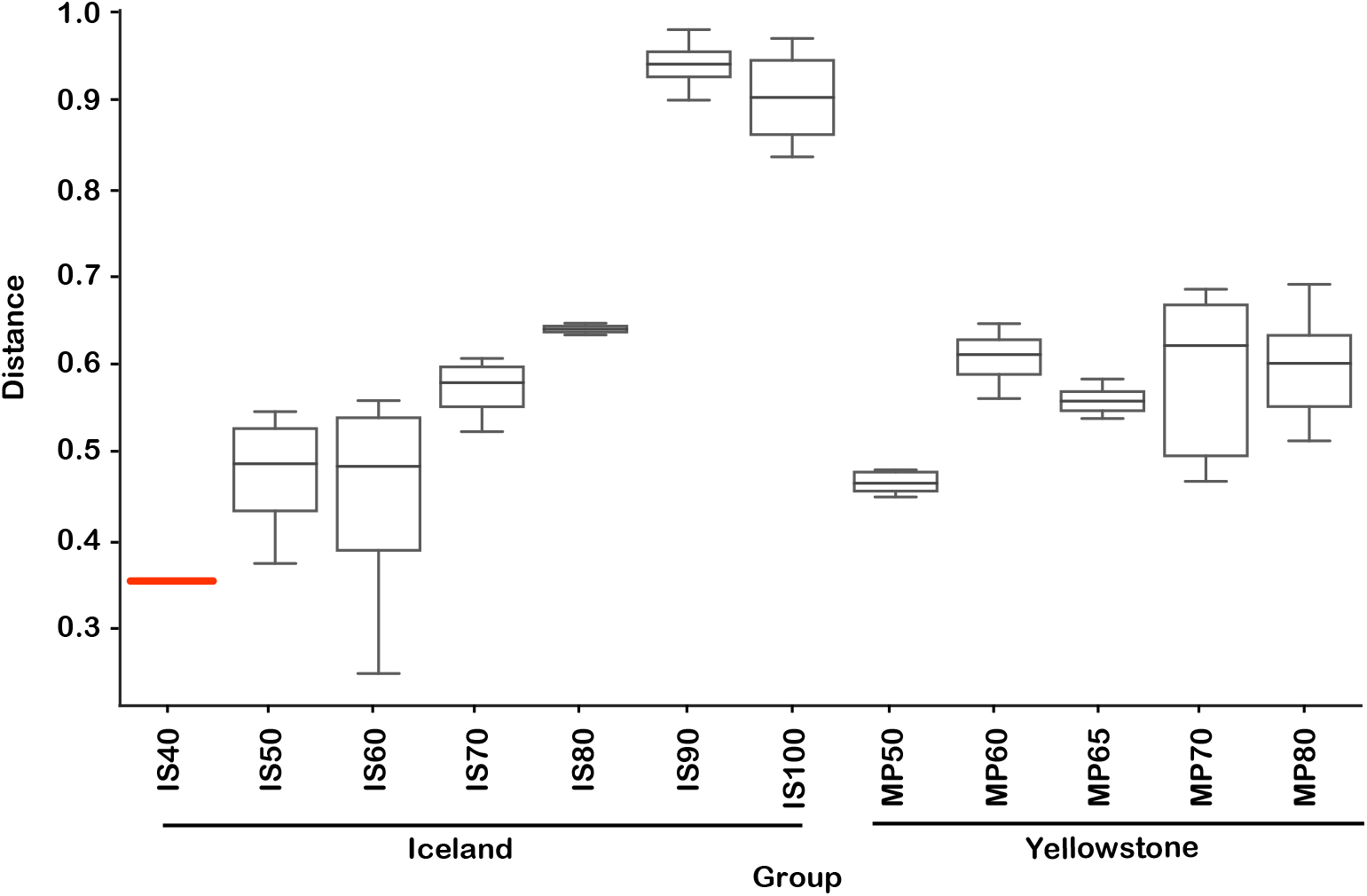
Weighted unifrac location-temperature group significance plot. Distances are relative to the lowest temperature group (Flúðir Hverahólmi, Iceland, 40°C). PERMANOVA F-test significance p=0.001. IS, Iceland.

### Microbial taxonomy across temperature gradients and hot springs

The 1729 unique sequence variants were assigned to 182 genera, corresponding to 5 phyla of Archaea (9 classes) and 40 phyla of Bacteria (86 classes) (**Figure 8 and 9**). The temperature gradient that forms in the three distinct hot spring systems creates distinct niches where organisms that are best adapted to those conditions thrive. In all those systems, such niches can be distinguished even macroscopically, based on the morphology and color of the mats (**Figures 1-3**). The deep amplicon sequencing that we achieved revealed the presence and relative abundance of numerous groups of organisms including the rare taxa. At the lowest temperature (38°C), the water and microbial mats of the Hverahólmi lagoon are dominated by a large diversity of mesophilic and mildly thermophilic heterotrophic as well as photosynthetic bacteria, including Alpha and Betaproteobacteria (*Roseomonas, Rhodobacter, Tepidimonas*), Bacteroidetes (*Chitinophaga, Saprospira*) and Cyanobacteria (*Cyanobium, Leptolyngbya*) (**Figure 9**). As the lagoon receives a constant stream of high temperature hot spring water, we also detected numerous thermophilic and hyperthermophilic bacteria and archaea in the lagoon (e.g. *Pyrobaculum*, Aquificae, Thermi) at <0.1% of total sequences. While the thermophilic organisms increase the alpha diversity, the lagoon being the most diverse of the sampled niches, they represent a physiologically inactive component of the community. Some, depending on their capacity to survive low temperatures and exposure to oxygen, may have the potential to colonize other hot springs by different dispersal mechanisms (water, wind).

**Figure 8.**
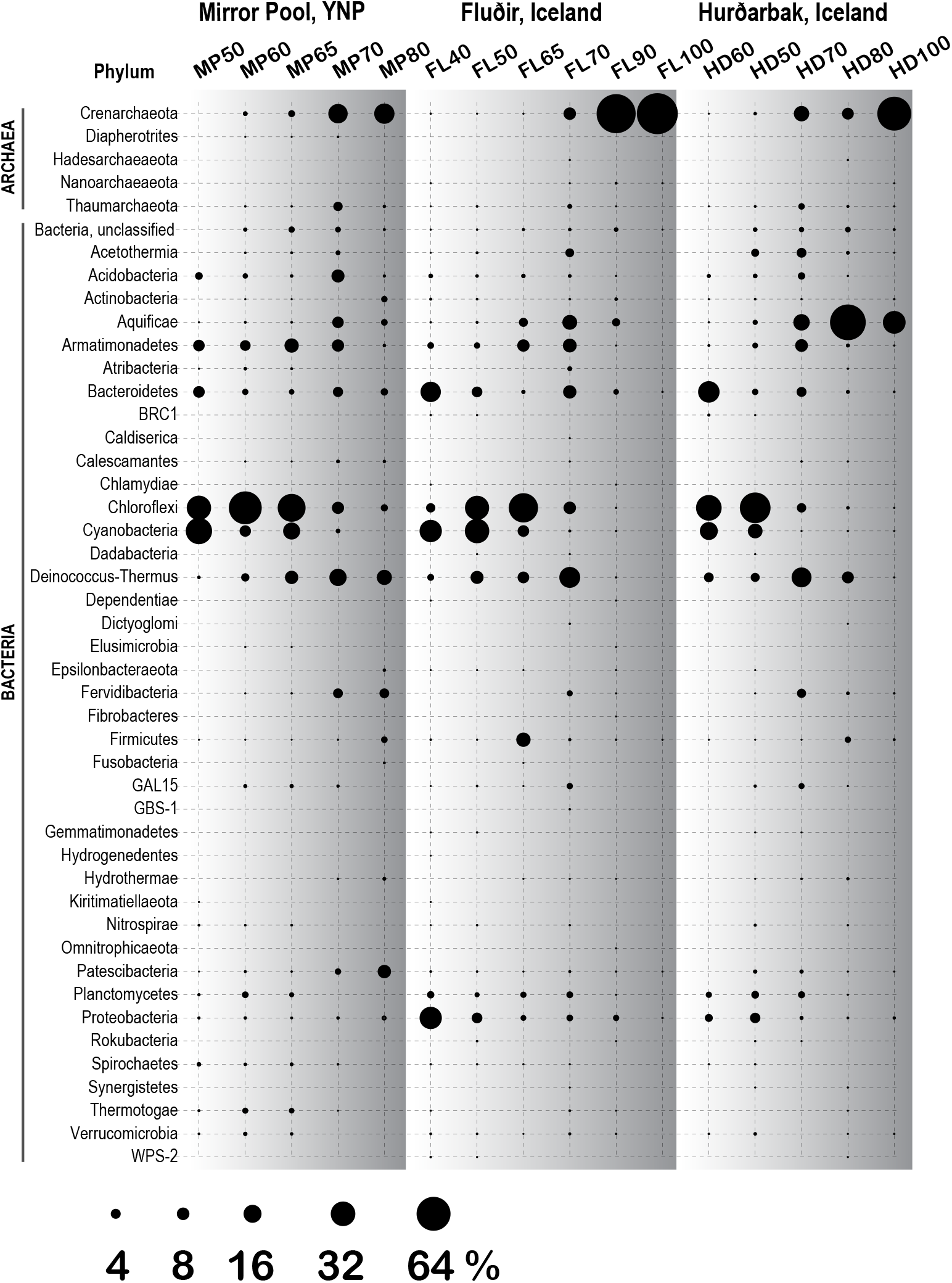
Bubble plot of microbial diversity (phylum level) at the three thermal springs depending on general temperature (40-100°C). Circle size indicates the inferred relative abundance, based on amplicon data (in %)

**Figure 9.**
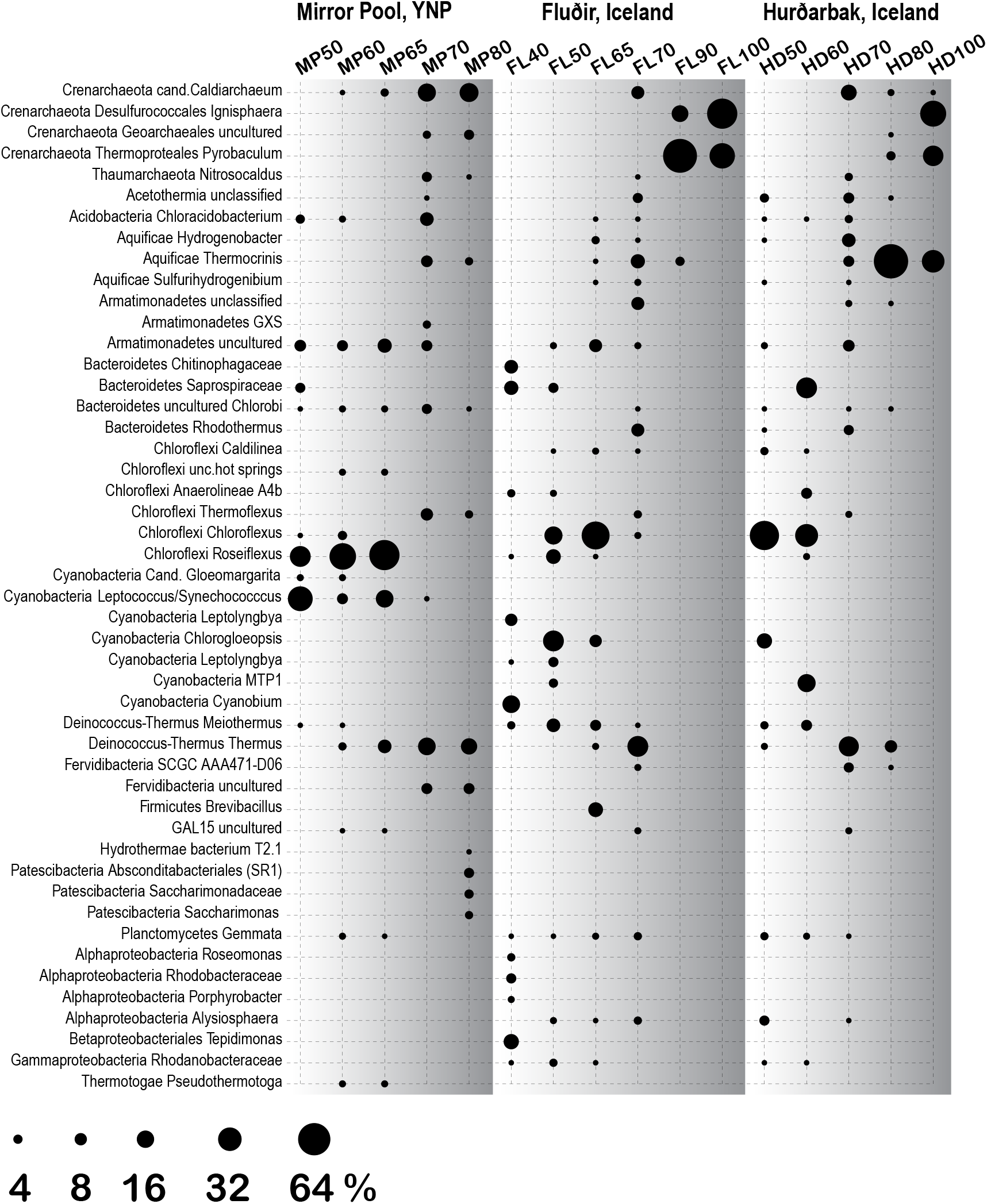
Bubble plot of most abundant genera/families at the three thermal springs depending on general temperature. Circle size indicates the inferred relative abundance based on amplicon data (in %)

The thermal areas (50-80°C) of all three hot springs share a variety of common organisms including Armatimonadetes, Chlorobi, Planctomycetes at the lower range, and Thaumarchaeota, Aquificaea (*Thermocrinis*) and Thermi, at the upper temperature range. There are though some specific differences, including dominance of *Caldiarchaeum* (Crenarchaeaota/”Aigarchaeaota”) in Mirror Pool, lineage that appears at lower abundance in the Icelandic springs. Aigarchaeaota are very common in alkalike springs from Yellowstone and were proposed to potentially chemoautotrophically use oxygen as a terminal electron acceptor [30]. Because the two Icelandic hot springs that we sampled have higher sulfate levels and are more reduced, the availability of oxygen may be limited in the higher temperature range and only sustain Aigarchaeaota in the flow stream as water cools and accumulates oxygen. We could not test redox levels in the different niches of the hot spring runoff therefore this remains hypothetical. The dominant community members between 50-65°C were oxygenic and anoxygenic chlorophototrophs (diverse Cyanobacteria, Choloroflexi and *Chloroacidobacterium*), forming characteristic green, orange and red mats depending on the site and temperature, as it has been shown in many other hot springs around the world (e.g.[28,31–33,29]). There were however some notable differences between Mirror Poll and the Icelandic features (**Figure 9**). *Synechococccus/Leptococcus*, an abundant member of the mats in Yellowstone, as well as *Gloeomargarita*, were absent in the two Icelandic hot springs. The striking absence of *Synechococcus* in Icelandic hot springs has been previously documented as long with an overall lower diversity of thermophilic Cyanobacteria [34,35]. Two potential explanations have been proposed. As the light levels in Iceland decrease significantly during the winter months, that could prevent colonization by some phototrophs. Alternatively, it has been hypothesized that thermophilic *Synecococcus* and other species that are sensitive to freezing and desiccation are less likely to survive the time required for dispersal from sources in North America or Eurasia [36]. As it has been documented that some Cyanobacteria are more sensitive to sulfide, that may also contribute to the differences between the sampled springs [4]. The Icelandic mats harbored however, diverse other Cyanobacteria, primarily at lower temperatures, including *Chlorogloeopsis* and *Leptolyngbya*. We also observed important differences across Chloroflexi between the compared two thermal regions. The Mirror Pool mats are dominated by *Roseiflexus*, while the Icelandic mats are primarily composed of *Chloroflexus*. There are potential complex physiological interactions between the prototrophic autotrophs and hetero-autotrophs in the mats, which may cause competitions between the various taxa [28]. At the same time, various Cyanobacteria and Chloroflexi may have different sensitivity not only to sulfide and redox levels but also to salt concentration and silica, another difference between the Yellowstone and Icelandic systems we studied here. The Icelandic mats may therefore represent alternative models to study interactions between these groups of organisms under distinct geochemical conditions.

Above 70-73°C, the upper temperature limit for photosynthesis, the microbial mats end sharply and all communities were composed primarily of Archaea, Aquificae, Armatimonadetes, Thermi and a variety of other phyla including many uncultured lineages (**Figure 8**). As the highest temperature in Mirror Spring was 82°C, we could not compare hyperthermophilic communities (>85 °C) between the Yellowstone and Iceland springs. Both sources of Hurðarbak and Vaðmálahver springs (90-100 °C) were dominated by the strict hyperthermophic Crenarchaeaota *Pyrobaculum* and *Ignisphaera* but also contained low levels of Nanoarchaeota. Hurðarbak also had a significant population of *Thermocrinis*, an autotrophic member of the Aquificae that dominated the community at 80C but was a minor component in both Vaðmálahver and Mirror Pool.

### Potential endemism at Icelandic and Yellowstone springs

Inspection of unique sequence variants revealed potential lineages that may be endemic to hot springs in Iceland or Yellowstone. Geographic separation of thermophilic communities or Archaea and Cyanobacteria has previously been shown to result in divergence of local populations that would ultimately lead to speciation [12,13,34]. While the goal of this research was not to specifically identify and study such organisms, phylogenetic analysis was used to characterize several of the closely related sequence variants that were abundant at one location but absent in the other. They included Archaea, Chloroflexi and Cyanobacteria (**Figure 10)**. The genetic distance between those sequences suggests they represent related but distinct species or subspecies. Some of the sequences from Mirror Pool are very closely related or identical to prior sequences reported from other Yellowstone springs. Because we did not find no close homologues from Icelandic hot springs in rRNA sequence databases, the analyzed Vaðmálahver and Hurðarbak amplicons had their closest relatives in hot springs from China. These preliminary observations may support the hypothesis that endemic species evolve in those hot springs. Nevertheless, the taxon differences could also be due to differences in chemical composition of the hot springs, which may have selected for related but globally distributed species. Future studies and analyses of similar hot springs in across the world would be required in order to better understand the links between the ecological diversity and evolutionary history of thermophilic organisms.

**Figure 10.**
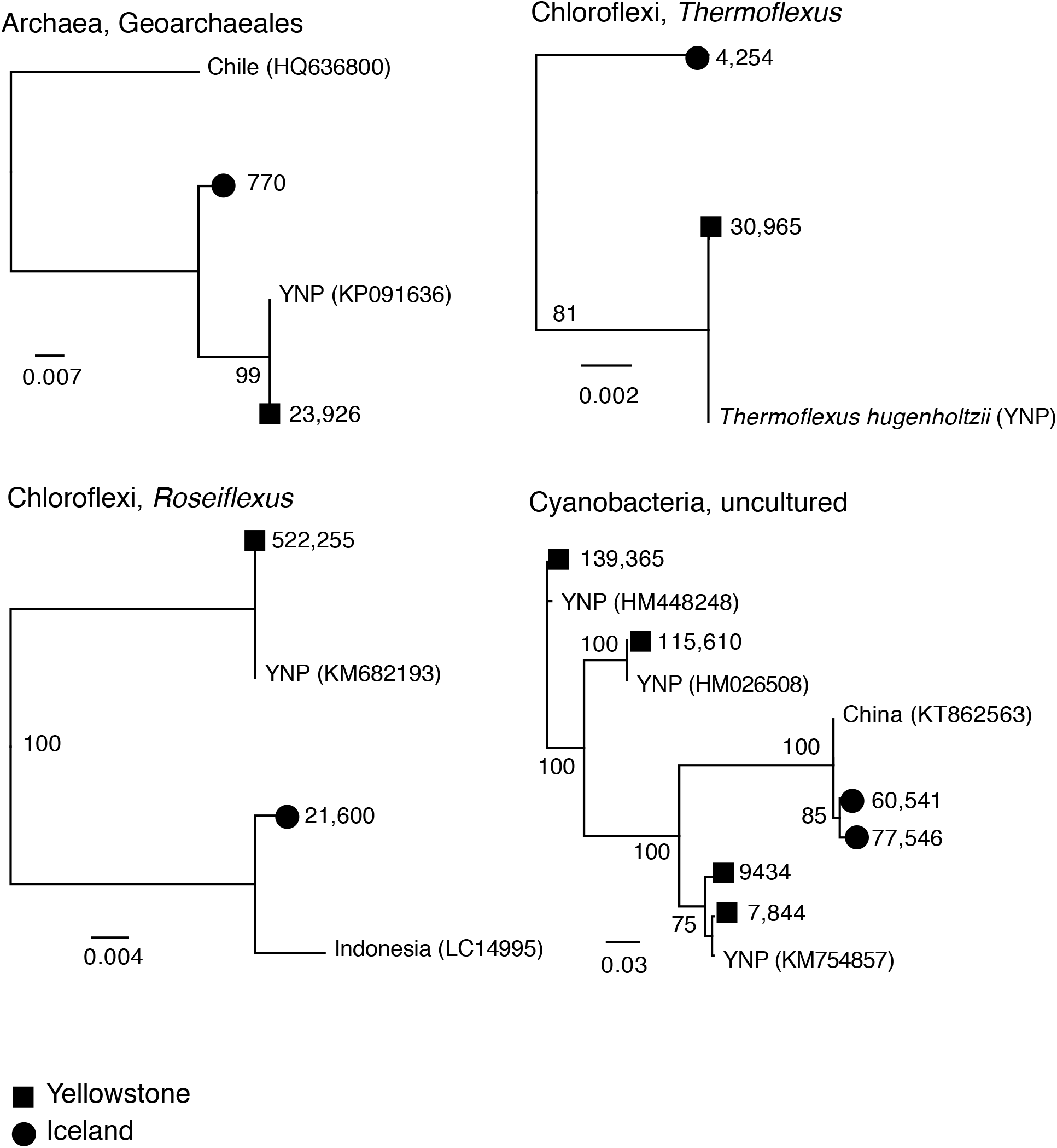
Neighbor joining trees (JC corrected distances) of endemic amplicon sequence variants (ASVs) from Iceland and Yelowstone (Mirror Pool). Numbers on branch tips indicate abundace across the entire dataset. Reference sequences and accession numbers from public rRNA databases and location were included. The numbers at nodes indicate bootstrap support. Scale bar indicates inferred number of substitutions per site.

## Conclusions

In this study we analyzed the microbial diversity across large temperature gradients in three hot springs from Yellowstone and Iceland that have not been previously investigated. One of the hypotheses tested was that the specific microbes that inhabit niches at different temperatures are similar across hot springs separated by thousands of kilometers. Also, we hypothesized that the overall microbial community structure is similar at similar temperature niches regardless of geographic isolation and small differences in chemistry. We identified an overall inverse relationship between the community diversity and temperature, with an abrupt decrease at the highest sampled temperatures (near boiling), where relatively few groups of organisms can survive. While there were specific differences at genus level between communities in Yellowstone and Iceland, potentially linked to differences in the chemistry of the hot springs, overall, they were remarkably similar, and temperature rather than geography site explained most of the microbial diversity.

## Acknowledgements

Environmental sampling in Iceland was under permits issued by Iceland’s National Energy Authority (Orkustofnun) to M. Podar and S. Björnsdóttir. We thank Dr. Jakob Kristjánsson for help with sampling and permits. Sampling in Yellowstone National Park was under permit YELL-SCI-5714 and we thank Stacey Gunther for help with sampling coordinating. We thank Adrian Gonzalez from the University of Tennessee Knoxville Water Quality Core Facility for chemical analysis of the water samples. This research was funded in part by a grant from the National Science Foundation (DEB1134877). Oak Ridge National Laboratory is managed by UT-Battelle, LLC, for the U.S. Department of Energy under contract DE-AC05-00OR22725.

## Author contributions

PTP and MP designed the study. PTP, SB and MP collected environmental samples. PTP and ZY performed DNA extraction, amplification and sequencing. PTP and MP analyzed the data and wrote the manuscript with input from SB and ZY.

